# EccDNA formation is dependent on MMEJ, repressed by c-NHEJ pathway, and stimulated by DNA double-strand break

**DOI:** 10.1101/2020.12.03.410480

**Authors:** Teressa Paulsen, Pumoli Malapati, Rebeka Eki, Tarek Abbas, Anindya Dutta

## Abstract

Extrachromosomal circular DNAs (eccDNA) are widespread in normal and cancer cells and are known to amplify oncogenic genes. However, the mechanisms that form eccDNA have never been fully elucidated due to the complex interactions of DNA repair pathways and lack of a method to quantify eccDNA abundance. Through the development of a sensitive and quantitative assay for eccDNA we show that the formation of eccDNA is through resection dependent repair of double-strand DNA breaks, especially micro-homology mediated end joining, and through mismatch repair. The most significant decreases in eccDNA levels occurred in cells lacking PARP1, POLQ, NBS1, RAD54, and FAN1. Further, a significant increase in eccDNA occurred in cells lacking c-NHEJ proteins DNA-PKcs, XRCC4, XLF, LIG4 and 53BP1. This suggests that when alt-NHEJ pathways are utilized to repair DNA breaks by necessity, the formation of eccDNA is increased. Induced and site-directed double-strand DNA breaks increase eccDNA formation, even from a single break. Additionally, we find that eccDNA levels accumulate as cells undergo replication in S-phase and that levels of eccDNA are decreased if DNA synthesis is prevented. Together, these results show that the bulk of eccDNA form by resection based alt-NHEJ pathways, especially during DNA replication and the repair of double-strand breaks.

## INTRODUCTION

Interest in eccDNA has been rapidly increasing because of the growing evidence of the prevalence of eccDNA cancer cells and their capability of promoting the genetic variability, tumorigenicity, and drug resistance.^1–9^ EccDNA increase the copy number of oncogenes in cancer, including full protein coding genes, microRNA, and novel si-like RNA.^1,10–12^ We have developed a method of quantifying eccDNA in cells lacking specific DNA repair proteins to show that the mechanisms by which eccDNA molecules are formed include the MMEJ, MMR and resection pathways. We also show that cellular eccDNA abundance is increased significantly when the classical-NHEJ (c-NHEJ) pathway is compromised.

Further, DNA damage, e.g. double-strand breaks (DSBs), in the genome promote eccDNA formation. This suggests that treatment of cancer cells with DSB inducing agents will increase eccDNA formation, which can promote gene amplification and drug resistance. We show that even a single DSB can induce eccDNAs in the vicinity of the break, suggesting that the role of DNA break is not simply to release linear DNA fragments that are circularized to form eccDNA.

We also show that during S-phase, when resection dependent DNA repair pathways are utilized and DSB like ends are created by replication fork stalling, eccDNA levels are significantly increased, most of them being lost as cells pass through mitosis when the nuclear envelope breaks down. Together, these results show that many eccDNA could be reduced through targeting the MMEJ/MMR/Resection proteins within a cell.

## RESULTS

### A QUANTITATIVE ASSAY FOR ECCDNA ABUNDANCE

To determine the abundance of eccDNA, we developed a quantitative assay that amplified sequences candidates representing the most abundant eccDNA molecules detected by next-generation sequencing of eccDNA. (EccDNA details in **Supplemental Table 1.**)

To eliminate chromosomal DNA with tandem duplications that may give a confounding signal, the eccDNA was enriched using a circular DNA plasmid isolation kit and contaminating linear DNA removed by digestion with an exonuclease for 24 hrs. We then carried out the inverse Q-PCR using outwardly directed primers that target only the junction sequence (**Supplemental Figure 1A)**. The abundance of mitochondrial DNA was shown to be unaffected by the various treatments and gene knock-outs used to test eccDNA formation, and this was used for normalization (**Supplemental Figure 1B-D)**. Further, the linear DNA was undetectable after the exonuclease digestion, thus removing any signal arising from chromosomal DNA (**Supplemental Figure 1E**).

### ECCDNA FORMATION IS INDUCED BY DOUBLE-STRAND DNA BREAKS (DSB)

We hypothesized that eccDNA formation was tied to DNA repair mechanisms, thus eccDNA levels would increase after DNA damage. Treatment of 293T cells with various agents that disrupt the DNA structure, most notably, those that induce double-strand breaks, increased eccDNA within 48 hours (**Figure 1B-N**). For example, the addition of cisplatin, which leads to DNA crosslinking,^13^ increased the abundance of eccDNA (**Figure 1B**). Induction of thymidine-dimers by UV radiation also increased eccDNA levels (**Figure 1C**). UV radiation induces thymine dimers and can also induce DSBs at high levels.^14^ Additionally, treatment with MMS, which leads to methylation of DNA causing replication stalling which can induce double-strand breaks,^15^ increases eccDNA abundance (**Figure 1D**). Finally, several compounds known to directly cause DSBs (Neocarzinostatin, Bleomycin, and X-rays)^16,17^ increased eccDNA abundance (**Figure 1E-G**). Interestingly, the increase of eccDNA by each type of damage plateaued around a 2-fold increase, suggesting that eccDNA production is limited either by repair dynamics, or by the amount of damage that can be withstood by a cell.

**Figure 1.**
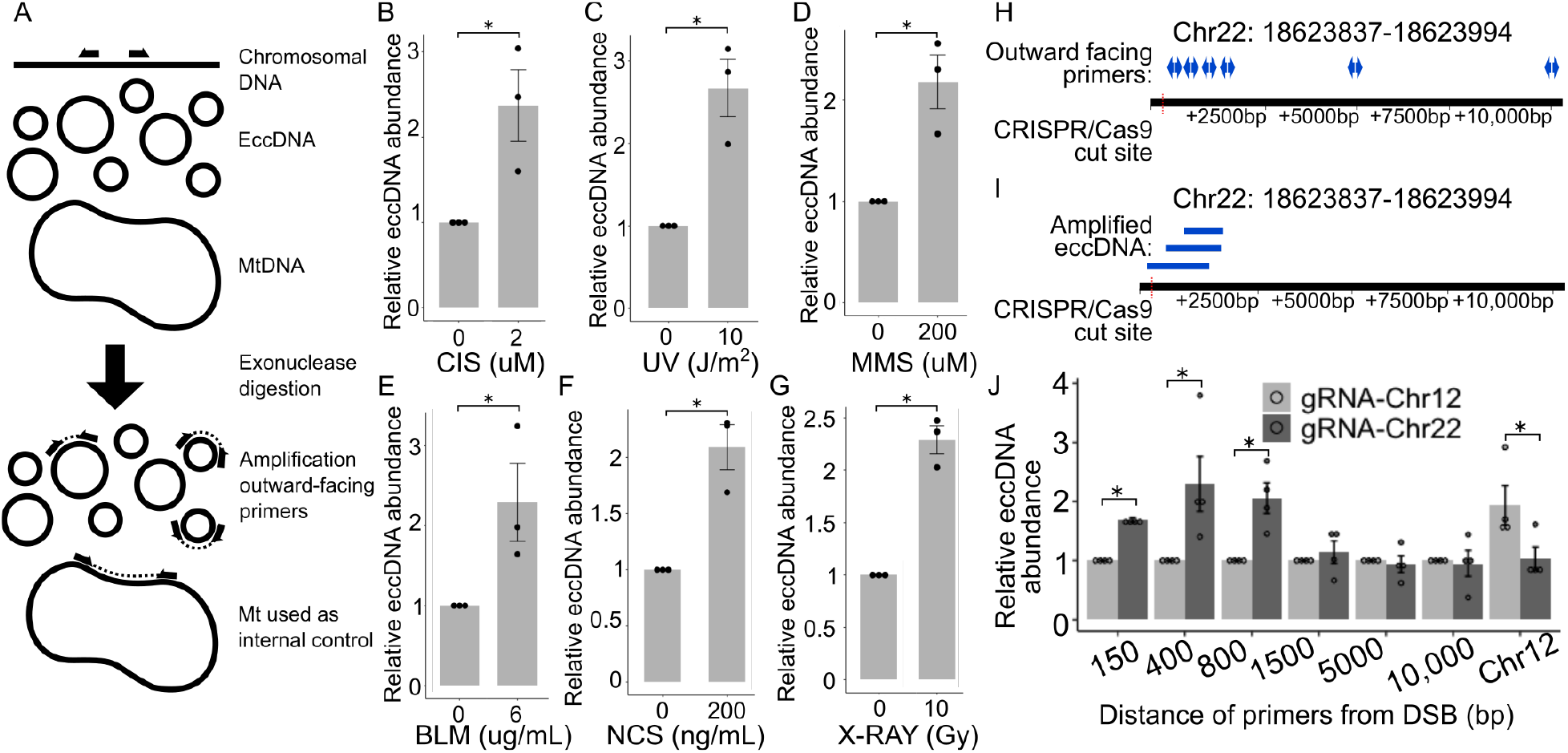
EccDNA formation is induced by disruptions to DNA structure. (A) Assay developed to quantify eccDNA. B-G: EccDNA levels measured by the QPCR assay in 293T cells 48 hours after the addition of (B) cisplatin (C) UV (D) MMS (E) BLM (F) NCS (G) X-rays. (H) Diagram of induced DSBs at locus Chr22:18624104 in 293A cells and the amplification of circles arising proximal to the cut site using outward facing primers (I) Diagram of eccDNA sequences amplified by the outward facing primers (J) Quantification of eccDNA isolated 48 hours after the transfection of an a p413 vector containing CAS9 and a gRNA sequence or a gRNA sequence targeting Chr22:18624104 or Chr12:117100086. P-values < 0.05 are indicated with a (*), < 0.01 (**), <0.001 (***).

### ECCDNA FORMATION IS INCREASED AT LOCUS OF DOUBLE-STRAND BREAK

Multiple DSBs on the same chromosome could release chromosomal fragments that are circularized to form eccDNAs. To determine whether a single DSB leads to the local induction of eccDNA, we induced a DSB at a specific locus in 293A cells by CRISPR-Cas9 mediated targeting of a specific location in the genome (Chr22:18624104). The DSB cut site on Chr 22 was chosen because this region naturally forms low levels of eccDNA which improves the sensitivity of the assay, and contains unique sequences which can be targeted by PCR primers. This site does not contain genic sequences and so eccDNA formation is not influenced by transcriptional changes, and its GC content is close to the genome average. Finally, the sgRNA sequence at Chr 22 had >99.9% specificity to the targeted site and its cutting efficiency was >60% (**Supplemental Figure 1F, G**). To ensure that the induction was specific to the area near the Chr 22 cut, we measured the eccDNA from an eccDNA hotspot on Chr 12 (**Supplemental Table 1**).

The amount of eccDNA produced from the Chr 22 locus was measured using outward-directed primer pairs located at distances of 150, 400, 800, 1500, 5000, 10000 bp from the DSB site (**Figure 1H, 1I**). EccDNA were stimulated proximal to the DSB: 150, 400, 800 bp away from DSB (**Figure 1J**). EccDNA further from DSB were not increased. (EccDNA listed in **Supplemental Table 1**). These results show that a DSB induces the formation of eccDNA from the chromosomal DNA proximal to the DSB. In contrast, the Chr 12 locus did not increase eccDNA production when cells were transfected with the Chr 22 sgRNA (**Figure 1J**) showing that the effect was local to the site of DSB cut. A similar locus-specific induction of eccDNA was seen with an sgRNA targeting an eccDNA hotspot on Chr 12 and primers ~300 bp away from the cut site. Therefore the induction of eccDNA near a targeted single DSB was not specific just to the Chr 22 locus.

### ECCDNA FORMATION IS SUPPRESSED BY c-NHEJ

To elucidate which DNA repair pathways form eccDNA as products of repair, we utilized our quantitative assay for eccDNA on an array of isogenic cell lines that have DNA repair genes knocked-out with CRISPR or inhibited by specific molecular inhibitors (**Figure 2A-R**). The results from human and chicken cells under normal growth conditions are summarized in **Figure 3C**. Human U2OS cells that lacked functional genes in the c-NHEJ pathway, *DNA-PKcs*, *XRCC4*, *XLF*, and *LIG4*, all had a significant increase of eccDNA; 53BP1, while not being a primary player in NHEJ, favors NHEJ by preventing hyper-resection at DSB ends, and loss of 53BP1 also increases eccDNA levels (**Figure 2B**). The critical role of DNA-PK in suppressing eccDNA production is further supported by the increase of eccDNA in human embryonic kidney derived 293T cells treated with the specific DNA-PK inhibitor (CAS-20357-25-9) (**Figure 2D**). Together, these data show that c-NHEJ suppresses eccDNA formation, and if compromised eccDNA levels rise significantly.

**Figure 2.**
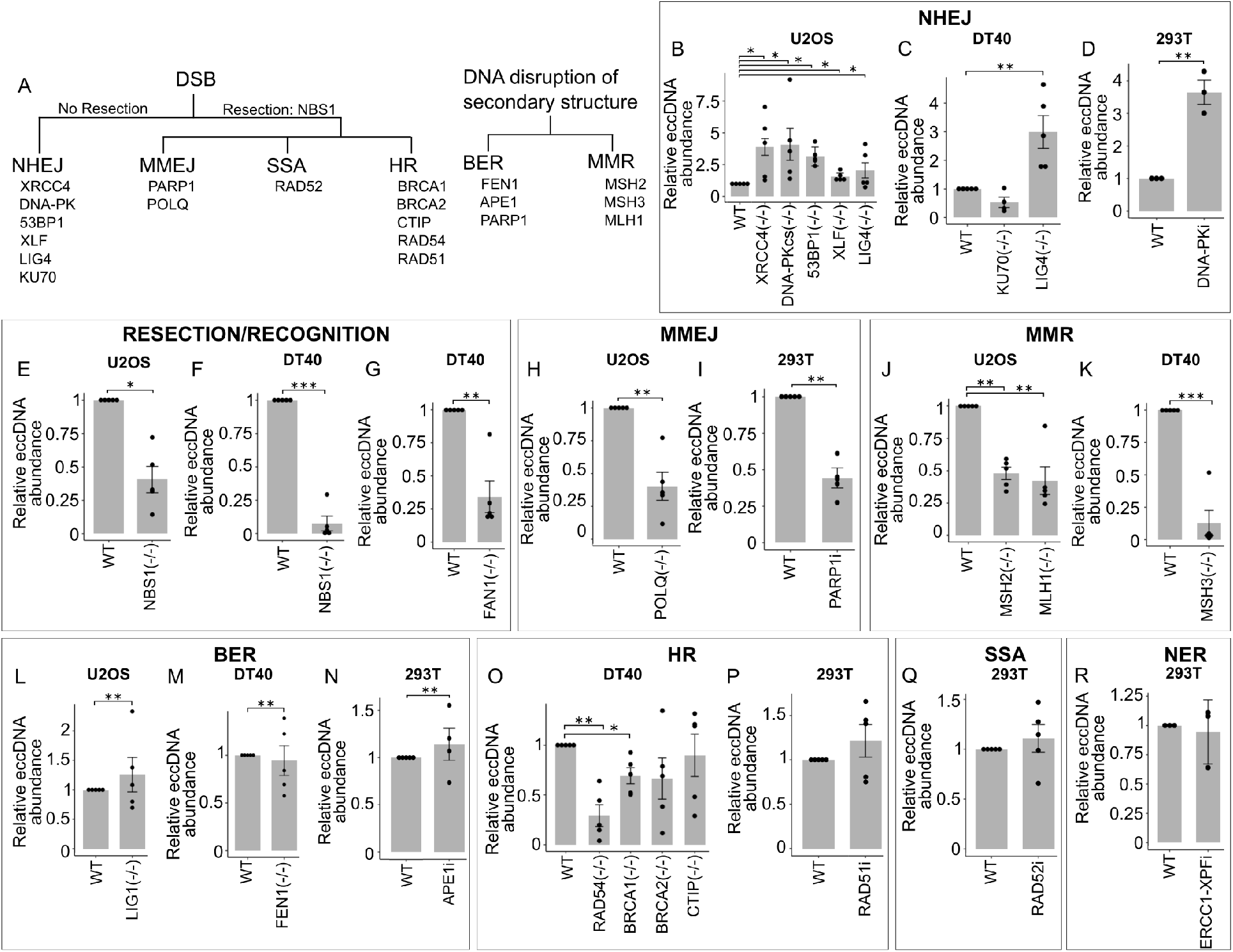
EccDNA formation is suppressed by c-NHEJ and increased by alt-NHEJ pathways. (A-E) Levels of eccDNA in U2OS cell lines with various knocked-out genes within DNA repair pathways. (A) Diagram of genes knocked-out or inhibited. Relative levels of eccDNA in different knock-out and small molecule inhibitor of DNA repair pathways: (B-D) NHEJ, (E-G) resection and recognition of DSBs, (H-I) MMEJ, (J-K) MMR, (L-N) BER, (O-P) HR, (Q) SSA, (R) NER.

**Figure 3.**
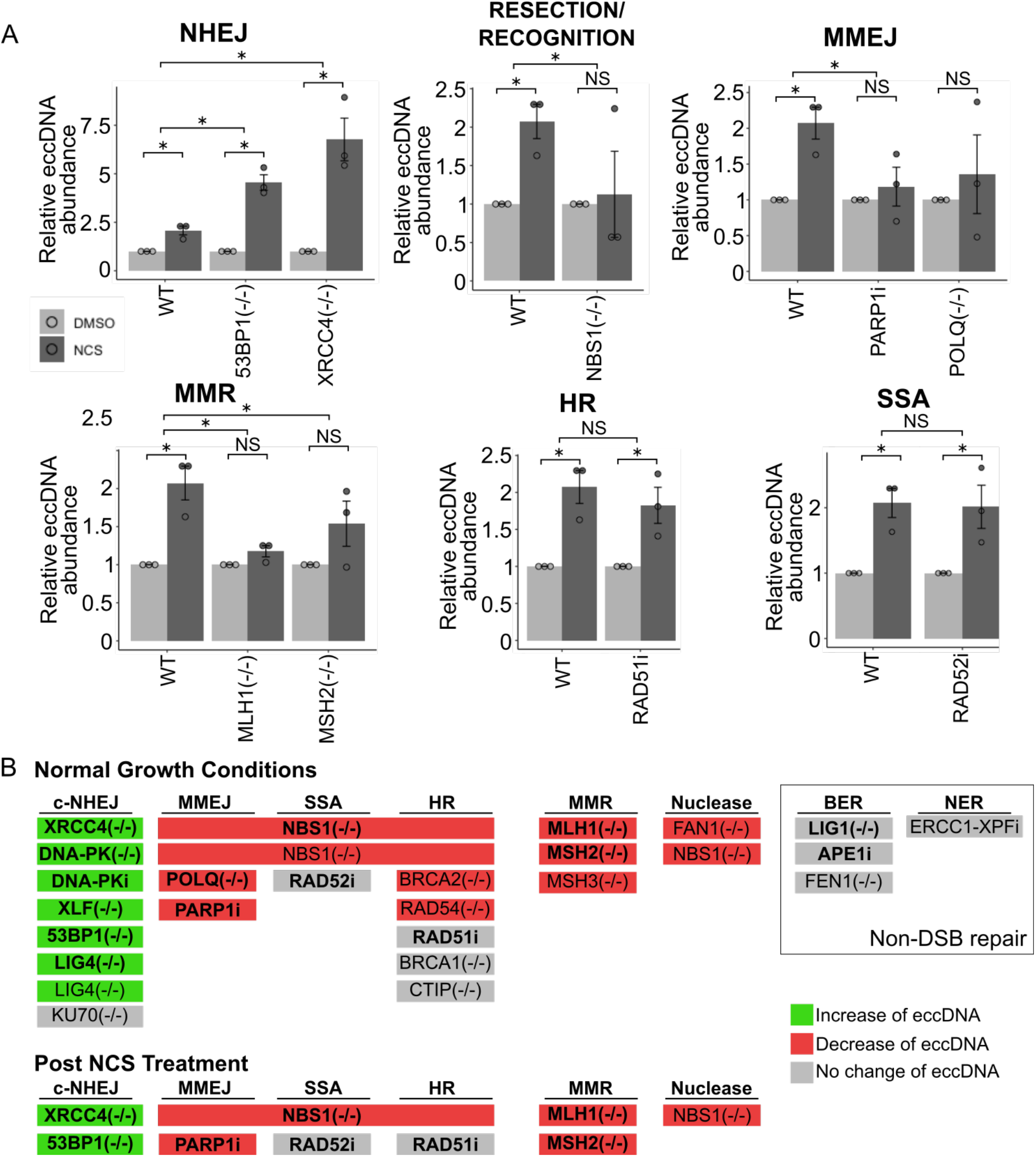
After DSB, cells lacking c-NHEJ have more eccDNA and cells lacking functional MMEJ and resection proteins have fewer eccDNA. (A, B) Levels of eccDNA after treatment of NCS (200 ng/mL) for 48 hours. (C) Graphical summary of whether mutants or inhibitors or a given repair pathway increased (green) or decreased (red).

To determine how general this result is, we used chicken DT40 lymphoma cells knocked out for various DNA repair genes. Lack of *LIG4*, important for c-NHEJ, in DT40 cells also increased the eccDNA levels (**Figure 2C**). The knock-out of *KU70* did not change eccDNA levels significantly, but we suspect that this could be explained by the higher levels of cell-death experienced in this cell line. KU70 is known to be an essential gene in mammalian cells,^18^ so that it is likely that the *KU70−/−* DT40 cells adapted to the loss of KU70 in some way that alters normal NHEJ DNA repair. Broadly, these results show that the formation of eccDNA is suppressed by the repair of the DSBs by the c-NHEJ pathway.

### ECCDNA FORMATION IS FACILITATED BY ALT-NHEJ REPAIR

We next tested eccDNA abundance in alt-NHEJ DNA repair pathways which require resection to rejoin DNA strands after damage, including MMEJ, SSA, and HR. NBS1, as a component of the MRN complex, is recruited early to DSB, and is critical for resecting DNA ends and a major factor in repair choice from c-NHEJ to end-resection dependent alt-NHEJ and HR repair.^19^ The knock-out of *NBS1* significantly decreased eccDNA abundance both in human U2OS and in chicken DT40 cells (**Figure 2E, 2F**). Further, loss of *FAN1,*^20,21^ a 5’-3’ nuclease implicated in interstrand cross-link repair and replication fork stability also significantly reduced eccDNA levels (**Figure 2G**). Together these data suggest that nucleases involved in DNA resection promote eccDNAs formation.

We next turned to the contribution of proteins within the MMEJ DNA repair pathway, which requires a small degree of resection and homology for repair.^22^ Loss of *POLQ*, a helicase-polymerase involved in unwinding DNA and facilitating the annealing of homologous ssDNA in MMEJ,^23^ significantly reduced levels of eccDNA (**Figure 2H**). PARP1 tethers DNA ends and interacts with XRCC1 and LIGIII to promote MMEJ,^24^ and the lack of PARP1 greatly reduces MMEJ.^22^ Addition of a PARP1 inhibitor, AZD2461, to 293T cells reduced levels of eccDNA (**Figure 2I**). The reduction in eccDNA upon inhibition of PARP1 and POLQ suggests that MMEJ is a major DNA repair pathway contributing to the observed endogenous levels of eccDNA.

The MMR pathway is known to have some interaction with MMEJ.^25,26^ We therefore tested whether the loss of proteins within MMR would alter eccDNA abundance. Knock-out of *MSH2* or *MLH1* in U2OS cells, as well as knock-out of *MSH3* in DT40 cells, led to a substantial decrease of eccDNA (**Figure 2J, 2K**). Combined, these data show that resection, MMEJ, and MMR all significantly contribute to eccDNA formation.

### ECCDNA FORMATION INDEPENDENT OF BER

PARP1 inhibition is known to disrupt base excision repair (BER) of some types of DNA lesions.^27^ To determine whether the PARP1 inhibitor is altering eccDNA production through the BER pathway, we tested eccDNA levels after the inhibition of two genes within the BER pathway. The knock-out of LIG1, the ligase utilized by BER,^28^ did not significantly change eccDNA abundance in U2OS cells (**Figure 2L**). Furthermore, the knock-out of FEN1, the endonuclease known to be necessary for BER by processing flap-containing intermediates^29^, did not change eccDNA abundance in DT40 cells (**Figure 2M**). The APE1 endonuclease is necessary at an early stage in BER because it is recruited to the apurinic sites to cut the DNA and recruit other BER proteins.^30^ Consistent with the lack of an effect of BER on eccDNA production, an APE1 inhibitor (Mx) did not significantly alter eccDNA abundance in 293T cells (**Figure 2N**). Therefore, the role of PARP1 in promoting eccDNA production is likely dependent on its role in MMEJ but not in BER. (Effectiveness of chemical inhibitors verified in **Supplemental Figure 2A-D**).

### ENDOGENOUS ECCDNA FORMATION MOSTLY INDEPENDENT OF HR PATHWAY

To test the contribution of HR to eccDNA formation, we tested cells lacking functional HR proteins including RAD54, BRCA1, BRCA2, or CTIP in DT40 cells (**Figure 2O**). Mixed effects were seen. RAD54 and BRCA1 loss significantly decreased eccDNA levels. BRCA2 loss may decrease eccDNA, but the variability of the result prevented us from reaching statistical significance. In contrast CtIP loss in DT40 or a RAD51 inhibitor on 293T cells (B02: **Fig. 2P**) did not decrease eccDNA levels. The mixed results make us wonder whether the effects seen with some of these genes are due to their roles in pathways not involving HR. For example, the loss of both BRCA1 and BRCA2 are known to reduce MMEJ activity and this could be why we see the decrease of eccDNA.^31^ Alternatively, BRCA1 is known to attenuate the c-NHEJ pathway and thus its loss may promote c-NHEJ and lower eccDNA formation.^32^ In addition, BRCA1 and BRCA2 may have an indirect role in endogenous eccDNA formation because of their activity in fork regression,^33^ or fork protection.^15,34^ The requirement of NBS1 from the MRN nuclease complex, but not of CTIP could be explained by the fact that the MRN complex has an endonuclease activity that is essential for end resection, that MRN can bind and resect DNA slowly but independently of CTIP and BRCA1^35^, or that other enzymes like DNA2 and EXO1 are also involved in strand resection with CTIP but after MRN action.^36^

In yeast the loss of MUS81, a resolvase used during the late stages of HR in D-loop resolution that is similar to RAD54, is required for eccDNA formation.^37^ The inhibition of RAD54 reduces strand invasion and reduces homology searching during HR.^38^ Thus, it is possible that RAD54 may have a role in the either the search of microhomology or even the strand invasion role we hypothesize in the model in **Fig. 4D** to stimulate endogenous eccDNA formation.

**Figure 4.**
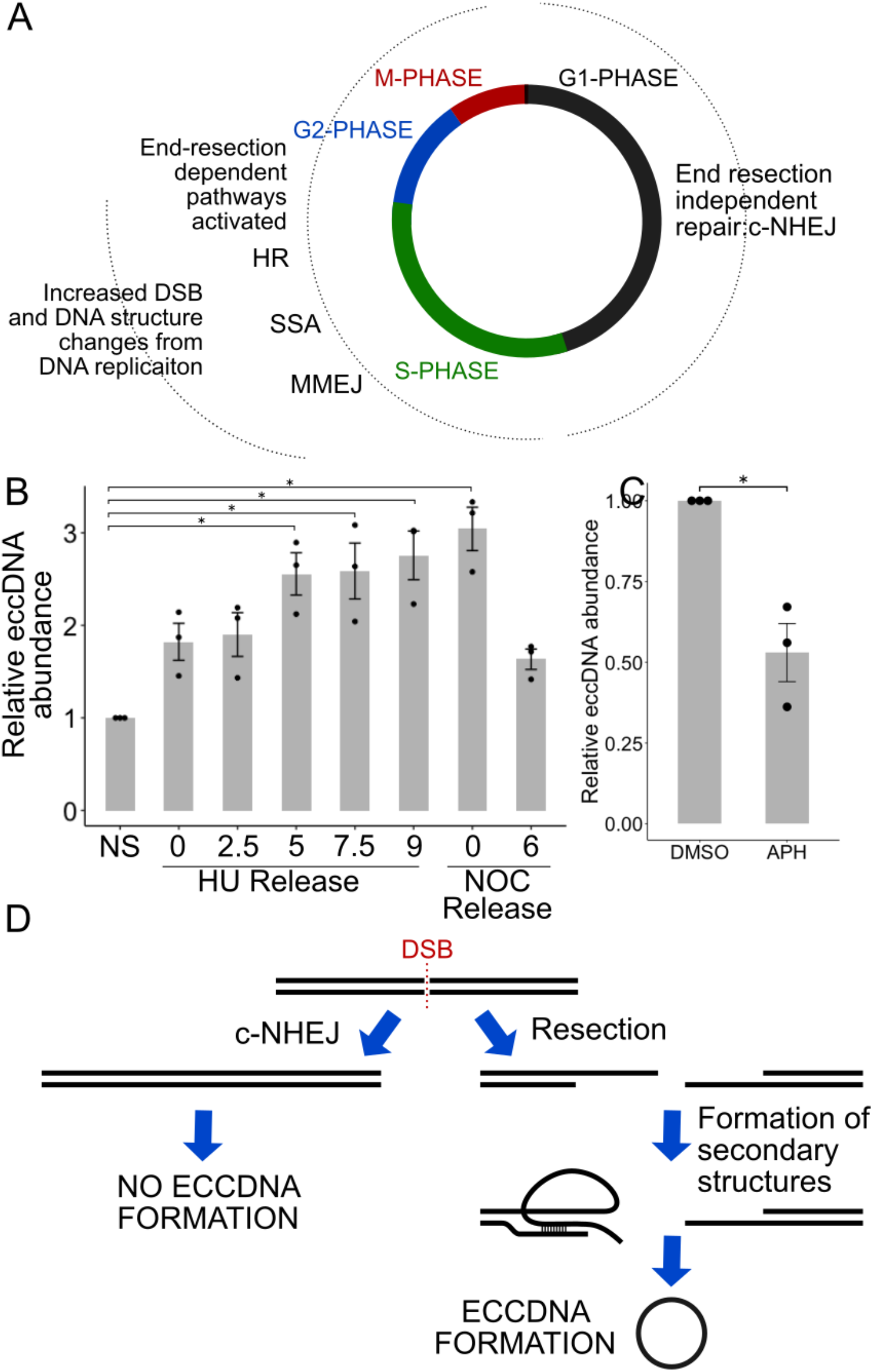
EccDNA formation is increased in S-phase, G2-phase, and M-phase of the cell cycle. (A) End-resection-independent pathway of repair is active in G1, while end-resection (and homology) dependent repair is more active in S and G2. (B) EccDNA levels in S-phase, G2-phase and M-phase. (C) EccDNA levels after the addition of 3 uM aphidicolin (APH). (D) Model of endogenous eccDNA formation from a single DSB.

### ECCDNA FORMATION INDEPENDENT OF SSA AND NER

To test other repair pathways downstream from MRN-mediated end-resection, we analyzed eccDNA levels in cells lacking functional proteins for single strand annealing (SSA) repair. RAD52 is necessary for SSA because it mediates strand invasion in a RAD51 independent manner.^39^ We found no change in eccDNA abundance after the addition of the RAD52 inhibitor, D-103, to 293T cells (**Figure 2Q**), suggesting that SSA is not essential for eccDNA formation. Further, the production of eccDNA is not dependent on the NER pathway, as tested with the inhibition of ERCC1-XPF with NSC16868 (**Figure 2R**).^40,41^

### CELLS LACKING C-NHEJ PRODUCE MORE ECCDNA AFTER DSBs

To examine the interaction between DSBs and the repair pathways implicated above in eccDNA formation, we induced DSBs in cells with compromised DSB repair by the addition of NCS) (**Figure 3C**). Cells lacking genes that promote the c-NHEJ pathway, *XRCC4* and *53BP1*, produced more eccDNA upon addition of NCS compared to WT cells (**Figure 3A**). This result is consistent with results above suggesting that c-NHEJ repair decreases endogenous eccDNA levels.

Further, cells compromised in alt-NHEJ pathways do not show an increase of eccDNA after DSB induction. The loss of PARP1 and POLQ (MMEJ); MLH1 and MLH2 (MMR), and NBS1 (resection) suppressed the increase of eccDNA after DSBs (**Figure 3B**), consistent with the hypothesis that alt-NHEJ, especially MMEJ, is important for eccDNA production. We conclude that alt-NHEJ and MMR promote eccDNA formation both under basal conditions as well as following the induction of DSBs by DSB-inducing agents (**Figure 3C**). The RAD51 or RAD52 inhibitors suppressed the production of eccDNA after NCS treatment (**Figure 3B**), but the decrease in the induction post NCS did not reach statistical significance from that in control cells (**Figure 3C**).

### ECCDNA FORMATION IS INCREASED IN S-, G2-, AND M-PHASE OF THE CELL CYCLE

The extent of endogenous DSBs and the utilization of different DNA repair pathways are different in different parts of the cell cycle (**Figure 4A, 4B**). DSBs are increased during normal DNA replication in S phase when the replication fork runs into nicks or other barriers to DNA replication.^42^ Further, c-NHEJ is used throughout the cell cycle, but repair pathways dependent on end-resection are used only in S- and G2-phase.^18^ Therefore, we hypothesized an increase of eccDNA through S-phase.

Cells blocked at the G1-S phase transition in hydroxyurea were released from the block by washing the cells, and cells harvested at 2.5, 5, 7.5, and 9 hours after release. FACS for analysis confirmed that the cells progressed through S-phase (**Supplemental Figure 3**). The eccDNA levels were significantly increased by the HU block, and increased further as the cells progressed through S-phase (**Figure 4B**). This suggests that DNA repair pathways that are utilized in S-phase, to repair DSB or resolve stalled or collapsed forks, etc. contribute to the formation of eccDNA.

We tested the levels of eccDNA in cells blocked in M-phase by nocodazole. The levels of eccDNA were elevated similar to the levels of the cells in the late S/G2-phase (**Figure 4B**). When we released the cells from M-phase for 6 hours to allow them to progress into G1, the eccDNA levels declined **(Figure 4B)**. This suggests that the eccDNA may be exposed to the cytoplasm during cell division and experience some degradation. Together, this shows that eccDNA formation increases during S-phase when DSBs appear and resection dependent DNA repair pathways are most active. Consistent with a role of DNA replication related DSBs in producing eccDNA, prevention of DNA replication with aphidicolin, an inhibitor of replicative DNA polymerases, lowers eccDNA levels (**Figure 4C**).

## DISCUSSION

EccDNA are formed by DNA repair pathways, especially after DSB and during replication (**Figure 4D**). Our results suggest that most endogenous eccDNA seen in normal cells and cancer cells are produced by a different process. The DNA repair proteins that are necessary for eccDNA after DNA damage are tied to MMEJ, DNA end-resection and homology searching, i.e. POLQ, PARP1, MSH2, MSH3, MSH6, MLH1, NBS1, FAN1, and RAD54.

Additionally, proteins that favor c-NHEJ, which suppresses resection based DSB repair, suppress the formation of eccDNA, i.e. XRCC4, XLF, DNA-PKcs, LIG4, and 53BP1. Together this shows that following even a single DSB, DNA repair pathways which resect the DNA and produce a single-strand tail promote eccDNAs formation. We hypothesize that the single-stranded DNA uses microhomology to form secondary structures with the double stranded DNA in *cist*, and this leads to aberrant DNA structures which are cleaved from the chromosome to form eccDNA (**Figure 4D**). The genetic information excised during the process could be restored by HR repair from the sister chromatid, so that not all eccDNA formation will accompanied by corresponding chromosomal deletions. This is consistent with a recent study on eccDNA containing the HXT6/7 gene under selection in yeast cells.^75^ We speculate that MMR may be involved in eccDNA formation because they have an undiscovered role in recognizing and repairing mismatches and looping structures that may occur during annealing of homologous sequences.^25,43^

The increase of eccDNA in hydroxyurea-blocked cells (when the cells have replication stress), and during S phase, suggests that endogenous eccDNA formation is connected to repair pathways associated with fork stalling. The decrease of eccDNA as cells pass through mitosis into G1 suggests that the eccDNA experiences some degradation in dividing cells.

It has been previously shown that the induction of two DNA DSBs induced by exogenous CRISPR/Cas9 induction within the same chromosome can lead to eccDNA formation from the excised DNA by c-NHEJ.^44^. These findings suggest that c-NHEJ can contribute to eccDNA formation, but only when there are two DSBs on the same chromosome. Such concordant DSBs *in cis* are unlikely to be common in normal cells, because that would lead to frequent chromosomal deletions, and so the endogenous eccDNA are most likely formed from single DSBs. However, this does not rule out eccDNA formation accompanied by chromosomal deletions in other contexts, e.g. when forming the large chimeric circles called ecDNA in cancer cells undergoing chromothripsis.

Recently, it has been shown in yeast that endogenous eccDNA formation is tied to SAE2 and MRE11, proteins that resect DNA in DSB repair.^37^ MRE11 of the MRN complex is important for eccDNA formation in yeast,^37^ consistent with our finding that NBS1 of the MRN complex is important for eccDNA formation in human and chicken cells. We could not use an MRE11 inhibitor Mirin to directly test the role of MRE11 in forming endogenous eccDNAs because of the very high toxicity of the drug at long intervals (**Supplemental Figure 2E**). MUS81 nuclease, which when paired with EME1 is involved in the unhooking of an interstrand cross-link, is also required for eccDNA formation in yeast.^45^ This is similar to the requirement we note for FAN1, though it is possible that MUS81 in yeast and FAN1 in human cells are required not to cut near a crosslink, but to cut a flap to release a circle in the model in **Figure 4D**.

The increase in eccDNA abundance following DSB could be relevant to the use of chemotherapeutic agents or radiotherapy, which lead to DSBs, and may, therefore, increase somatically mosaic eccDNA, thus increasing the genetic variation of cancer cells and potentially leading to adaptation of the cancer to therapy. Compromise of the c-NHEJ pathway will lead to increased abundance of eccDNA. This could be relevant to cancers where c-NHEJ is known to be down regulated in cancers such as breast cancer^46^ as it suggests that the loss of c-NHEJ can increase genomic instability and tumor adaptation through increased formation of somatically mosaic eccDNA.

## METHODS

### ECCDNA QUANTIFICATION

The eccDNA was isolated from the various knock-out cell lines and treated cells using a HI Speed midi-prep DNA isolation kit. The linear DNA was then digested using ATP-dependent plasmid safe DNase. The remaining circular DNA was purified using DNA Clean & Concentrator-5 Kit. Then QPCR was performed using SYBR master mix.

### CELL CULTURE

HeLa cells were cultured using standard techniques. The effectiveness of the drug was confirmed using the techniques described in published literature (**Supplemental Figure 2**).

### FACS ANALYSIS

Cells were fixed in 70% ethanol, washed, and stained with propidium iodide.

## ACKNOWLEDGEMENTS

This work was supported by the NIH grant R01 CA060499. We thank members of the Dutta and Abbas labs for suggestions and helpful discussions.

## AUTHOR CONTRIBUTIONS

Teressa Paulsen wrote the manuscript and performed the experiments. Pumoli Malapati contributed biological replicates in 293T cells. Rebeka Eki and Dr. Tarek Abbas generated and characterized the U2OS KO cell lines. Dr. Anindya Dutta managed the project and edited the manuscript.

