## Supplemental Materials for "EccDNA formation is dependent on MMEJ, repressed by c-NHEJ pathway, and stimulated by DNA double-strand break"

**DNA repair of double-strand breaks by end-resection and homology dependent repair promote eccDNA formation: SUPPLEMENTAL MATERIALS**

Teressa Paulsen, Pumoli Malapati, Rebeka Eki, Tarek Abbas, Anindya Dutta

**Supplemental Figure 1:**

**(A) Outward facing primers target the known junction sequence of the eccDNA** and will therefore specifically amplify the sequence only if it has become circularized. The primers are designed to create amplicons of similar size though the eccDNA targeted by the primers range from 180-1300 base pairs.

**(B-D)The mitochondrial DNA levels between samples are not significantly changed** by the genotoxic agents used or by DNA repair gene mutations in (B) 293T cells treated with the indicated genotoxic agents, (C) U2OS KO cells, (D) DT40 KO cells.

**(E) The purification of eccDNA** through exonuclease digestion removes contaminating genomic DNA, as measured by primers for the U6 gene, but preserves circular DNA like mtDNA, as measured by primers targeting the mtDNA. Cells were treated with DMSO or NCS.

**(F) Cutting efficiency of CRISPR/Cas9 system at Chr22 assessed by mismatch created by repair of cut.** T7 endonuclease digestion of PCR amplicon after de- and re-naturation will cut mismatched duplexes. Amplicons were from chr22:18623440-18624398 after transfection of plasmid carrying CAS9/sgRNA targeting either chr22: 18624104 or chr12:117100086.

**(G)** Percentage~60% of DNA amplicons were cut by T7 endonuclease, indicating a successful cut by CRISPR/Cas9 followed by erroneous repair by NHEJ.

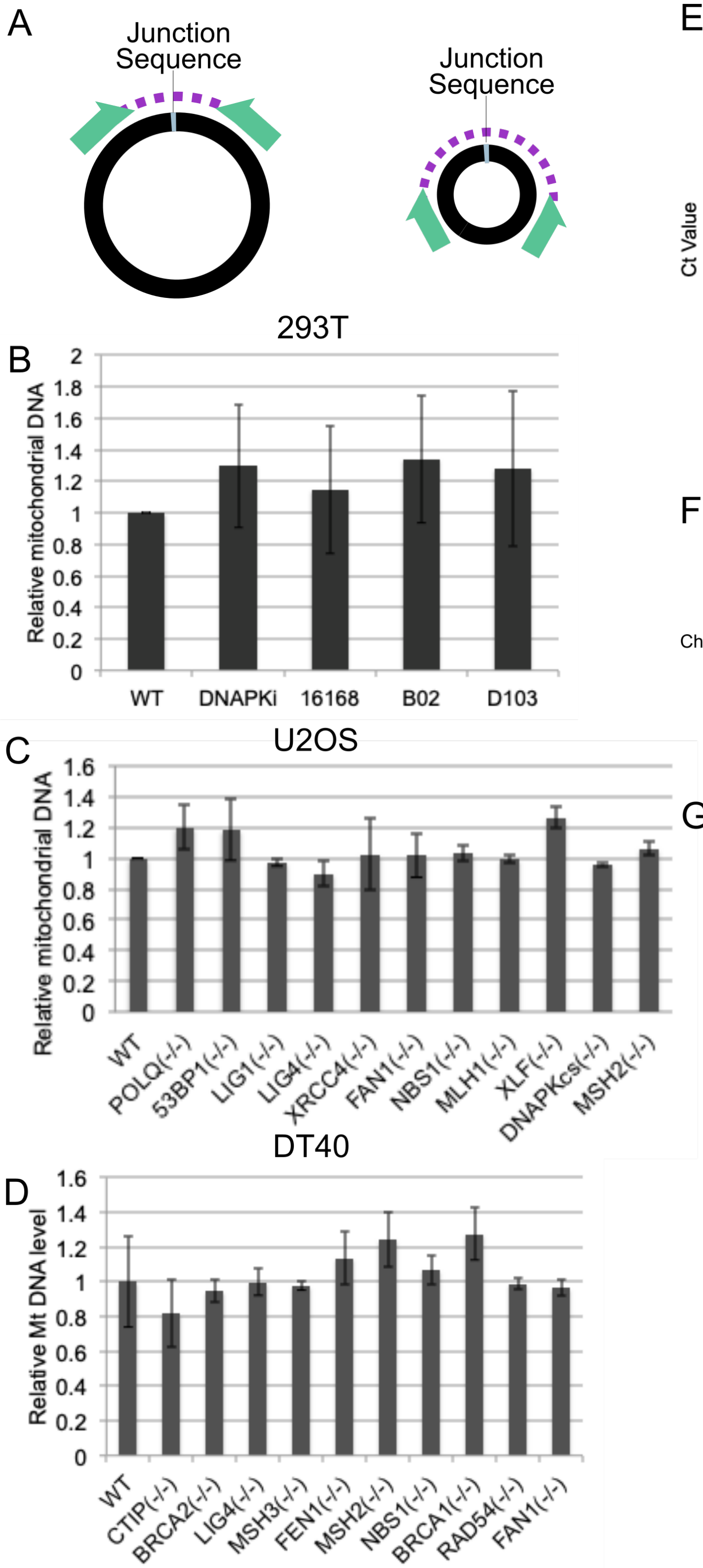


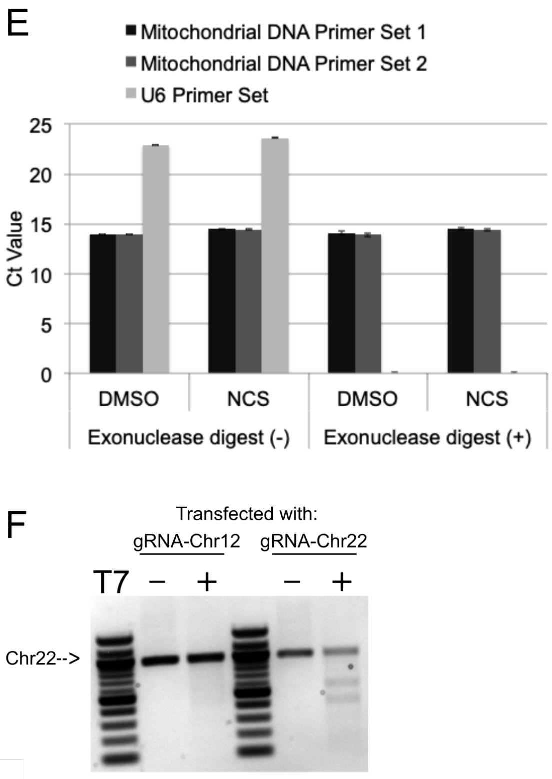


G

Efficiency of Chr22 cut

| | Area |
|---|---|
| Uncut | 56331 |
| Cut band 1 | 43466 |
| Cut band 2 | 37274 |
| Percent Cleaved | 58.90 |

**Supplemental Figure 2:**
**(A-D) The efficiency of small molecule inhibitors utilized in study.** Treatment of 293T cells decreased cell viability as expected from functional inhibitors. (A) Cisplatin (CIS) and NSC16168 (inhibitor of ERCC1-XPF), (B) D-103 (Rad52 inhibitor) or B02 (RAD51 inhibitor), (C) cisplatin and B02, (D) Mx (APE1 inhibitor in the BER pathway) with CrVI (hexavalent chromium).
**(E) Mirin significantly reduces cell survival during 48 hour treatment.**This indicates that treatment with Mirin for 48 hours is not useful for eccDNA measurement because of the death which occurs in ~90% of cells.

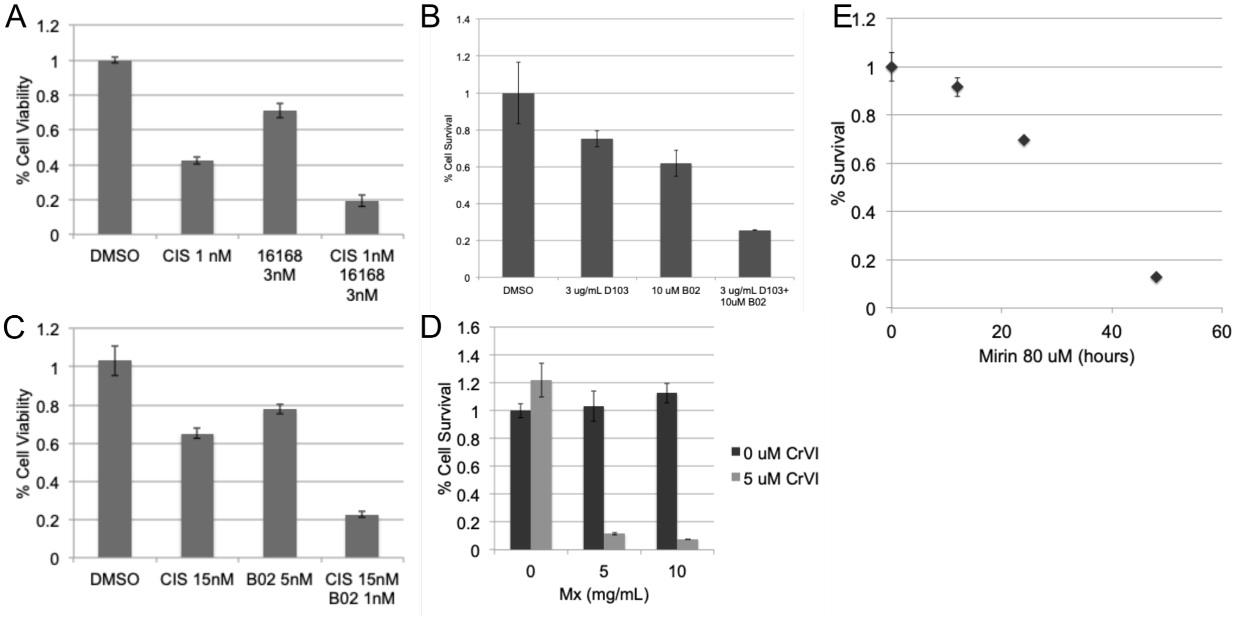

**Supplemental Figure 3: Propidium Iodide FACS profiles to show cell-cycle profile of HeLa cells.** To ensure that cells were progressing through S-phase as predicted, we analyzed the cells at several points after release from hydroxyurea block: 0, 1.5, 3, 4.5, 6, 7.5, and 9 hours. We also confirmed that cells were stalling in M phase after the addition of nocodazole.

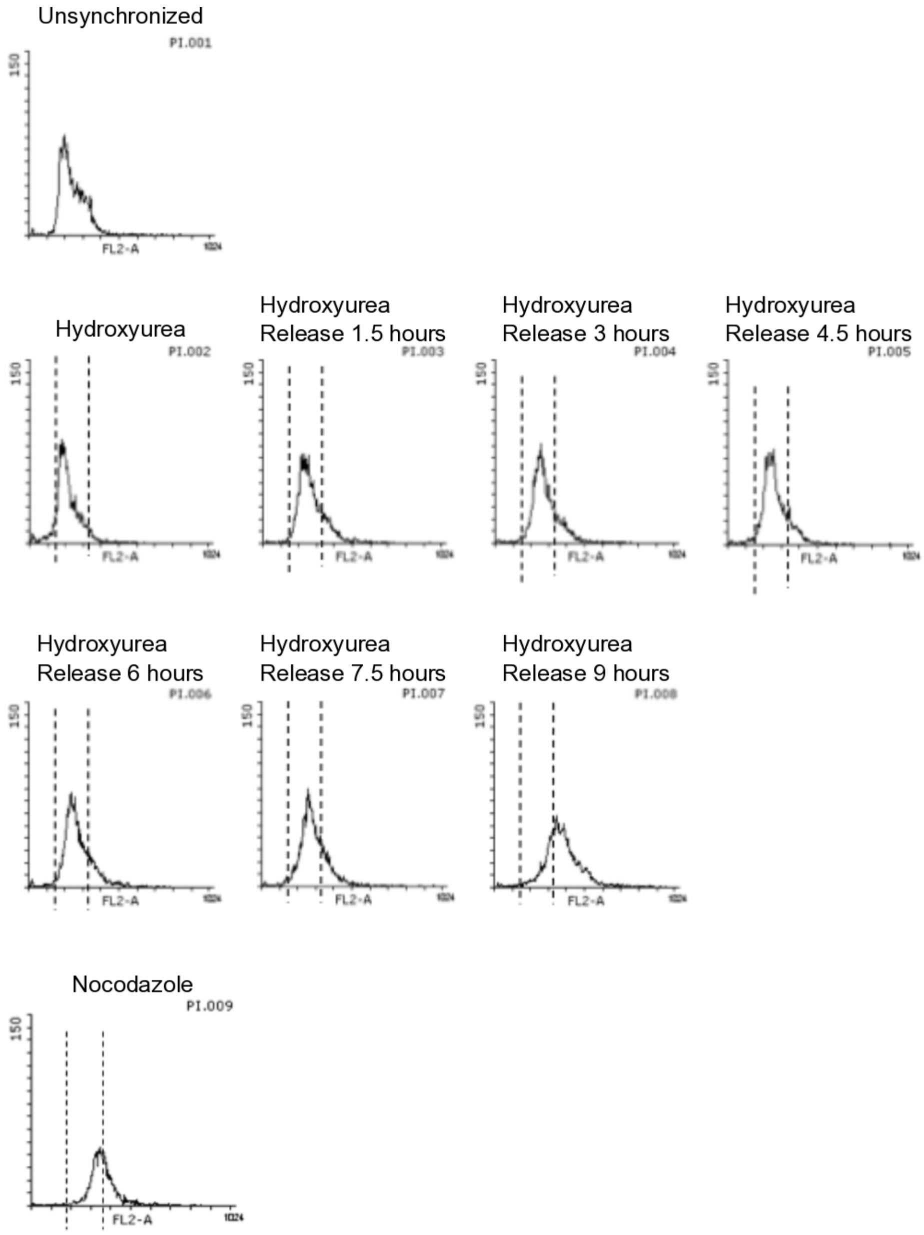

**Supplemental Table 1: Chromosomal loci interrogated by inverse PCR,** selected because they produced highly abundant eccDNA in human or chicken cells as detected in eccDNA libraries prepared by rolling circle amplification and high throughput sequencing from past research.[7] The abundance was listed as the number of times the junction sequence was sequenced, thus giving an estimate of the percentage of the specific eccDNA within the entire population of eccDNA. Sequences of outward facing primers, size of inverse PCR amplicon and sequence of the amplicon with the two parts that are joined to form the circle are given. The two parts are indicated by bold and regular font; the junction is at the shift between bold and regular font. UF: Upstream facing, DF: Downstream facing.

**A)** Human eccDNA. **B)** Chicken eccDNA. Some primers amplified more than one eccDNA, therefore some primers have more than one sequenced amplicon.

A)

| **Human OF Primers and Amplicon Sequences** | | | | | | | | | | |
|---|---|---|---|---|---|---|---|---|---|---|
| **#** | **UF Primer sequence** | **DF Primer Sequence** | **Coordinates of eccDNA by NGS** | | | **Abundance by NGS** | **Size of circle from NGS (bp)** | **Coordinates of sequence in genome** | **Size of Amplicon (bp)** | **Sequence of OF primer PCR amplicon** |
| 1 | GAGATAGCCAAGCTCCCATG | TACTACCATTGCAGATTGCG | chr1 | 245530219 | 245530472 | 265246.5 | 253 | chr1:245530078-245530685 | 567 | **GAGATAGCCAAGCTCCCATGTTTATTGCAGCACTATTCACAATAGTCAAGAGTTGGAAGCAATCTAAGCATCCATCAATAGACAAATGAATAAAGAAAAGGTGGTACATATACAGAATGGAGTACTCTTCAGCCATAAAGAAGAATGAGATCCTGCCATTTGCAACAACATGGATGGAACTGGAGGTCATTATGTTAAGTGAAATTAGCGAGGCCCAGAAAAACAGACTTCACATGTTTTTATGTGTGCTGCCCAGATGACACAATTCAGGAAAATCTTTCAGCTCCTACCAGAGAGTAAAGCTCTTCAGCTGTAGTGG**CTCAACATCATTCATTGATCATCATAGAAATGCAAATCAAAACTACAATGAGATTATCATCTCACCACAGTTAAAATGGCTTTTATCCAAAAATCAGGCAATAACAAATGCTGGTGAGGATGTGGAGAAAAGGGAACCCTCATACACTGTGGATGGGAATGTAAATTAGTACAACCACTGTAGAGAACAGTTTGGAGCTTCCTCAAAAAACTAACAATTGAACTACCACGCAATCTGCAATGGTAGTA |
| 2 | CCTCTGAAAGTGCTGGGATTACAG | GATCACGAGGTCAAGAGATGGAG | chr16 | 83,620,081 | 83,620,204 | 130172 | 123 | chr16:83620011-83620259 | 121 | **GATCACGAGGTCAAGAGATGGAGACCATTCTGGCCAACATGGTGAAACCCCGTCTCTACTAAAAGTACAAAAATTAGCTGGGCGTGGTGGTGC**TCGCCTGTAATCCCAGCACTTTCAGAGG |
| 3 | GGCCCTCTGGTGGACATCTCATTAA | CTGGTGGGCGCTATGGAATCTT | chr12 | 117100431 | 117100599 | 8358.2 | 168 | chr12:117100381-117100661 | 251 | **CTGGTGGGCGCTATGGAATCTTGTCCCCTGGAGAGGCACACAGCCAGGGCAGAACATCAAGGTCAAGGCTCTCCTGAAGGCTCTGCAGTGCTTAGTGACACCACCATCAATGACCGTCAGGTATCAGGTCTGTTCTGAACGTTAGTCT**TCTCCATCCAGTTGGGGTGACTGGGGCCGATGCTGTTGCCTGTGTAATGTGATTCCTCCTCCTTAAAATAAGGATAAGTTAATGAGATGTCCACCAGAGGGCC |
| 4 | ACCACCCTGG[illegible] | TGAACTGAA[illegible] | chr6 | 43484330 | 43484598 | 10.1 | 268 | chr6:43488800-43490545 | 270 | **ACCACCCTGGCCACAGAGGTGGGCCTGGCCTTTTCCCAAGCCTAGGATCAAGATTCACGCAGACAGGAAACACTTCCAATAATTGCCTACACACTGGGCTCCCCATGTTGCTCTGACTACAGCCCAAGAGCTCAAATAGGCCAAGGCCTGACTATGAAGA**TGATCCTAACTTCTTATCCTAGGGTGAGAGTAACAGGGACACTATCCAACCTTCCCTTCTTGATAAAGGGAGGAAATGATCTGATCTAGGAGAAAAGTCCCTTCAGTTCA |
| 5 | CGCTCCCAGAGCTGATAGGA | GGGGTTCTTCCAAGTGGCTCT | chr14 | 69,346,610 | 69,347,101 | 7352.5 | 491 | Chr14: 69346689-69347123 | 336 | **GGGGTTCTTCCAAGTGGCTCTGGGACCCTCTGCCCTGGAGGCTGGGCCTGAGACATAGAGGATGGGGGAGCCATGGGCGGAGCTCAGGCTGGAAAAACAGAGCCCCTCAGTCCAGAAAGAGGAAAGCCTCAGGAGAGCCTGGGGCCAAAATGCATGCTCA**CCAGCTGTGTTCTGGCCCTGAGATTCTAAGCAGTTCACCCCCTGCTTAATGATGCCGGGGAGTGCTTCCTTCACGGGGACCCTGAGAGAAAAGGCAGCAGTCAGGCTTGTGCTTTCCCCCCCAAGGCTTACCTACACCCTCTGCCAGACGTCAGCATCCTATCAGCTCTGGGAGCG |
| 6 | GCCTCCCAAGTAGGTGGGAT | GAGAATTGCTTGAACCCTGGAG | chr11 | 100,650,466 | 100,650,750 | 2880 | 284 | Chr11: 100650414-100650783 | 361 | **GAGAATTGCTTGAACCCTGGAGGTGGAGGTTGCAGTGAGCTGAGATCGCGCCACTGCAATCCAGCCTGGGCGGCAGAGTGAGACTCTGCCTCAAAAAAAAAAAGAGCAAAGTCAAGTCCATTGTCTTATACGGT**ACTTGCACATTTTTTAATGACTTCAACTTTCTTAAGAGTCAAGTCGGCCGGGTGTGGTGGCTCATGCCTGTAATCCCAGCACTTTGGGAGGCCAAGGCAGGCAGATTACTTGAGGTCAGGAGTTTGAAACCAGCCTGGCCAACATGGTGAAACCCCGTCGCTACTAAAAATACAAAAATTAGGTGGGCATGGTGGCATGCACCTATAATCCCACCTACTTGGGAGGC |
| 7 | TACTACCATTGCTAGATTGC | GAGATAGCCAAGCTCCCAT | chr1 | 245530571 | 245531755 | 247.9 | 1,184 | chr1:245530303-245530648 | 306 | **TACTACCATTGCTAGATTGCGTGGT**GAAAGATTTTCCTGAATTGTGTCATCTGGGCAGCACACATAAAAACATGTGAAGTCTGTTTTTCTGGGCCTCGCTAATTTCACTTAACATAATGACCTCCAGTTCCATCCATGTTGTTGCAAATGGCAGGATCTCATTCTTCTTTATGGCTGAAGAGTACTCCATTCTGTATATGTACCACCTTTTCTTTATTCATTTGTCTATTGATGGATGCTTAGATTGCTTCCAACTCTTGACTATTGTGAATAGTGCTGCAATAAACATGGGAGCTTGGCTATCTC |
| 8 | GATCACGAGGTCAAGAGATGGAG | CCTCTGAAAGTGCTGGGATTACA | chr19 | 49093402 | 49093582 | 6.9 | 180 | chr19:49091220-49091256 | 122 | **GATCACGAGGTCAAGAGATGGAGACCATCCTGGCTAACACGGTGAAACCCCATCTCTACTAAAAATACAAAAAA**TTAGCCGGGCGTGGTGGCGAGCGTCTGTAATCCCAGCACTTTCAGAGG |
| Mitochondrial DNA F:GATCAGGACATCCCGATGGTG R:AGGCGCTTTGTGAAGTAGGC | | | | | | | | | | |

B)

**DT40 OF Primers and Amplicon Sequences**

| # | UF Primer sequence | DF Primer Sequence | Coordinates of eccDNA by NGS | | | Abundance by NGS | Size of circle from NGS (bp) | Coordinates of sequence in genome | Size of Amplicon (bp) | Sequence of OF primer PCR amplicon |
|---|---|---|---|---|---|---|---|---|---|---|
| 1 | ATGCAAACCTGCCTTCGTTC | GCAGTGGTATGGGACCTGTC | chr11 | 2300736 | 2300935 | 19280.6 | 199 | Chr11: 2300736 - 2300969 | 200 | **GCAGTGGTATGGGACCTGTCACAGGAACCATATGGTCGTCTGCTTCTTCTGCAAAGCAGCCGTCACAAGAGAGGAGGATGCTGCA**GGGCTGCATGAATAAGCAGTGAGGAGCAAACCCACAAGGAAATTCAGGCAGTGCTTGGTTTTGCATAACAGTGCTATTTTGTCTGAGTTGTGGTTTACTGTTATGTTGTACAAGAAGAGACTATCGGATGCAAAGCCTCAACTCTGTAACTGGGAGGAGAGAGGGACAGCCATTACATCCATCAGTGAAACTGGAGGAGATGATGAACGAAGGCAGGTTTGCAT |
| 1 | ATGCAAACCTGCCTTCGTTC | GCAGTGGTATGGGACCTGTC | chr11 | 2300736 | 2300935 | 19280.6 | 199 | Chr11: 2300736 - 2300969 | 330 | **GCAGTGGTATGGGACCTGTCACAGGAACCATATGGTCGTCTGCTTCTTCTGCAAAGCAGCCGTCACAAGAGAGGAGGATGCTGCATATTGCTGCTGCCTCTGGAACGACGATATTTAGGT**CATCCATCAGTGAAACTGGAGGAGATGATGAACGAAGGCAGGTTTGCAT |
| 2 | TAAGCCTCAATGTGTGTTCAGCT | GAGGCAATTCTGAAAGAAGCCG | chr4 | 1460467 | 1460815 | 8535 | 348 | Chr4: 1460467 - 1462710 | 174 | **AAGCCTCAATGTGTGTTCAGCTGGAATGCTGTCTAGCTATACATCCCTTATTAATGCCATATTAACAGACACAGCTGTTCAATATGCAGTATGCAAACACACATGGATGTAAACAATCAGAGCAGAGGAGAGCTTTTGTTTCCTCGGCTTCTTTCAGAATTGCCTCCTCACTGTAAGCCTCAATGTGTGTTCAGCT**GGAATGCTGTCTAGCTATACATCCCTTATTAATGCCATATTAACAGACACAGCTGTTCAATATGCAGTATGCAAACACACATGGATGTAAACAATCAGAGCAGAGGAGAGCTTTTGTTTCCTCGGCTTCTTTCAGAATTGCCTC |
| 2 | TAAGCCTCAATGTGTGTTCAGCT | GAGGCAATTCTGAAAGAAGCCG | chr4 | 1460467 | 1460815 | 8535 | 348 | Chr4: 1460467 - 1462710 | 180 | **TAAGCCTCAATGTGTGTTCAGCTGGAATGCTGTCTAGCTATACATCCCTTATTAATGCCATATTAACAGACACAGCTGTTCAATATGCAGTATGCAAACACACATGGATGTAAACAATCAGAGCAGAGGAGAGCTTTTGTTTCCTCGGCTTCTTTCAGAATTGCCTCCTCACTGTAAGCCTCAATGTGTGTTCAGCT**GGAATGCTGTCTAGCTATACATCCCTTATTAATGCCATATTAACAGACACAGCTGTTCAATATGCAGTATGCAAACACACATGGATGTAAACAATCAGAGCAGAGGAGAGCTTTTGTTTCCTCGGCTTCTTTCAGAATTGCCTC |
| 3 | AGGAGCATCAACACTGATGCA | CCCCTAGATGGTCTGGACCT | chr3 | 25576742 | 25577047 | 5161.2 | 305 | Chr3: 25576742 - 25577047 | 205 | **CGTCTTGCTATAGCCCCCATCACCAACACACCTCTCACATTCCTCAAAATGCACCTTGAGACCAGAAACGAGCCCCCCTTGTTACCACGAAGCCATAACAAGCCCACAGCAACCACGTGCTGTCAGAGGCAGTGCCCAGGAACCCCAACCATGTCTCCCCAGCCTT**TGTCACTTGAAAGCTCACTGCATCAGTGTTGATGCTCCT |
| 3 | AGGAGCATCAACACTGATGCA | CCCCTAGATGGTCTGGACCT | chr3 | 25576742 | 25577047 | 5161.2 | 305 | Chr3: 25576742 - 25577047 | 170 | **CCCCTAGATGGTCTGGACCTTTATGCATAGACCTGATGCAGTGCTTTTTCAAATGCCTGAAATTGATAAAAGGATTCTGGTGGATTTTGGCCCCAAGCAGGCAGATGCCTCCTGTATACACAGTGTGCTGTACAGCTTCTACTGCATCTC**ACTGCATCAGTGTTGATGCTCCT |
| 4 | CTGCACTGTGATCAGCAGCT | GGAGGACAGATGGCTGCAC | chr4 | 82442526 | 82442718 | 5096.5 | 192 | Chr4: 82442326 - 82443624 | 177 | **GGAGGACAGATGGCTGCACCGCAGTCACTGCCAGCTGACACCTGCACAGGT**TGAGCTCTCAGAGCTCCCCGTCACCATTTCTTGCCTTCTGATCAGCAGTGTGCCTGTGAGTGAGAACTCACACGCAGCCCTTTTCAGGGCCCACGGGAGATGGAGCTGCTGATCACAGTGCAG |
| 4 | CTGCACTGTGATCAGCAGCT | GGAGGACAGATGGCTGCAC | chr4 | 82442526 | 82442718 | 5096.5 | 192 | Chr4: 82442326 - 82443624 | 179 | **GGAGGACAGATGGCTGCACCGCAGTCACTGCCAGCTGACACCTGCACAGGTTGAGCT**CTCAGAGCTCCCCGTCACCATTTCTTGCCTTCTGATCAGCAGTGTGCCTGTGAGTGAGAACTCACACGCAGCCCTTTTCAGGGCCCACGGGAGATGGAGCTGCTGATCACAGTGCAGCTCTC |
| 4 | CTGCACTGTGATCAGCAGCT | GGAGGACAGATGGCTGCAC | chr4 | 82442526 | 82442718 | 5096.5 | 192 | Chr4: 82442326 - 82443624 | 303 | **TGGCTGCACCCTCCCTGTCCGACTCATCCTCTTCAGACAGCCCCGAGTCAGC**TTCGGTGGGAGCTGGATCTGGTGGGGCAGCCTTGACGGGGGGCTCAGGCTCAGCTGAGTCCAAGCTGGGCTCCACTATGTGGAGCTCGGTGCTGTCCAGCTCTGCCTCTCCTTCCTCCTCCTCCTCTTCCGAGCTCTCAGCCTCTTGCTGAGCCACAGGACCCCCCGGGCATGGAGCCCCCTCTGCTCCCCAGCT |
| 5 | TTAGTTTCTGCTCGGCAGCA | TCAAACTCCGAGCACCAGTC | chr2 | 69137247 | 69137609 | 1057.7 | 362 | Chr2: 69137047-69137904 | 360 | **TCAAACTCCGAGCACCAGTCGTCTCGCTGTCAGGCAGTGCATGCTGGCTGCCTGCCTTATGGTGTGTGG**AGCACCAAGGCAGACAGTAGTTCAGAGCACTTTGCTGTTGTAAAATAAGGTTTCAAGCTCAGAGACACTGGTTTGCGCTCTCAAAGATGCAGATTCTGATCATCATCACATTTCATCTCGGTGATCTGTATCACTGAGTCCCAACAGTATGTAAGTTTGCATCAGATTCTCCCCTAAGTTCACTTCTTGCTTAGTTCACTGTTATGCTGCAGGTCTTTTTTTTTTAACCCTTCCATTGAGGGAGGTAAATGTAAAATGCTCAAGTGACTGCTGCCGAGCAGAAACTAA |
| 6 | ACTTCATCCAGTCCAGTGACAG | GAGGCTTCCACAACCTTTCAGT | chr8 | 13132838 | 13132985 | 1014.2 | 147 | Chr8: 13132654 - 13133032 | 335 | **ACTTCATCCAGTCCAGTGACAGCAGTACATCAGCAGATCCATACTGGCGTATCCTGCTGGTT**CTTCACACCTGTAGGTTGCCAGTTTTAGTCTGATCAGCCTAAAATGTTTTTTTTCAGTTGCACATGTGCTTTAGGACAGGTGTTGTACTGAAAGGTTGTGGAAGCCTC |
| 7 | GGGACAGGCTGAGAGCAGAG | GGCTTGGGAAGCAGTCATGC | chr2 | 20383499 | 20383673 | 1014.2 | 174 | Chr2: 20383321 - 20383744 | 199 | **GGCTTGGGAAGCAGTCATGCACATTG*GCC*TACCTGGCATGGGATTTGAACATCTTCACCCACAGTGTAAGACAGTGTGCATTGTGATGTGTAACTCCCATTGTTTTTTCTGCTATTTGCAGTGTAACGTGTGGATTAGTGCCCAAGTCATCTCTGCTCTGCTCTCAGCCTGTCCCTAATGGCTTGGGAAGCAGTCATGCACATTG**GCCTCAGCTGCTCTGGTTCCTACCTGGCATGGGATTTGAACATCTTCACCCACAGTGTAAGACAGTGTGCATTGTGATGTGTAACTCCCATTGTTTTTTCTGCTATTTGCAGTGTAACGTGTGGATTAGTGCCCAAGTCATCTCTGCTCTGCTCTCAGCCTGTCCC |
| 7 | GGGACAGGCTGAGAGCAGAG | GGCTTGGGAAGCAGTCATGC | chr2 | 20383499 | 20383673 | 894.1 | 174 | Chr2: 20383321 - 20383744 | 379 | **GCTTGGGAAGCAGTCATGCACATTGGCCAATTCACAGGCTGAAGAAGTGCAGAGGGCAGTACTGGGCACTTAACATTGCATTCTGTTAGAGTAATTTTTATCAGCGCGCTGTAAGTACACAATTTGGCATAACTAGTGCCTGGCGCTGCCTGATCCCTCACTCAGCTGTCAGT**CCATCTCCATCTCTGACCAGACAGGTGGTTGGCTGCCTCAGCTGCTCTGGTTCCTACCTGGCATGGGATTTGAACATCTTCACCCACAGTGTAAGACAGTGTGCATTGTGATGTGTAACTCCCATTGTTTTTTCTGCTATTTGCAGTGTAACGTGTGGATTAGTGCCCAAGTCATCTCTGCTCTGCTCTCAGCCTGTCCCTAATGGCTTGGGAAGCAGTC |
| 7 | GGGACAGGCTGAGAGCAGAG | GGCTTGGGAAGCAGTCATGC | chr2 | 20383499 | 20383673 | 894.1 | 174 | Chr2: 20383321 - 20383744 | 375 | **GGCTTGGGAAGCAGTCATGCACATTGGCCAATTCACAGGCTGAAGAAGTGCAGAGGGCAGTACTGGGCACTTAACATTGCAT**TCTGTTAGAGTAATTTTTATCAGCGCGTTGTAAGTACACAATTTGGCATAACTAGTGCCTGGCGCCTCATGTCACAAACGCTCTCAGCCACAGCCATCTCCATCTCTGACCAGACAGGTGGTTGGCTGCCTCAGCTGCTCTGGTTCCTACCTGGCATGGGATTTGAACATCTTCACCCACAGTGTAAGACAGTGTGCATTGTGATGTGTAACTCCCATTGTTTTTTCTGCTATTTGCAGTGTAACGTGTGGATTAGTGCCCAAGTCATCTCTGCTCTGCTCTCAGCCTGTCCC |
| 8 | CACGACTGACAGGATTATCAGGC | AAGGATTTGTGCATGCTTCTGAC | chr1 | 30701201 | 30701303 | 894.1 | 102 | Chr1: 30701201 - 30701303 | 190 | **CACGACTGACAGGATTATCAGGCCCACAGTATCGCTCCTATCCCTTTTTCAGCAATGCCGCTTTCCAGGTTATTCCCTCACAGCGAATACCTGTTTT**ACAAAGGCTATATACCATACTGTAAACTGAGGCCTGTTCTCCCTTGTCAGAAGCATGCACAAATCCTTTTTATATGAAGGGGAGTTGGGTGCA |
| Mitochodrial DNA F:ACCCCCAACTCAAACACTCG R:GGTGGTTCCTAAGACCAACGG | | | | | | | | | | |

**Supplemental Table 2: Characteristics of the eccDNA candidates in human cancer cell lines and DT40 cells. (A**) Human cancer cell line eccDNA candidates **(B)** DT40 cell line eccDNA candidates. (microhomology between eccDNA sequence and flanking genomic sequence indicated by arrows)

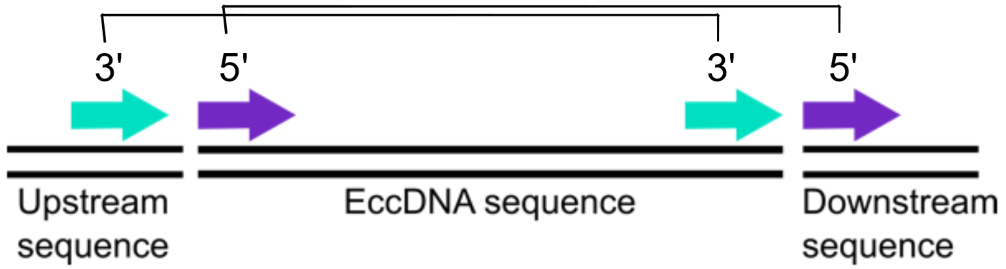


A

| Human Cancer Cell Line EccDNA Candidates | | | | | | |
|---|---|---|---|---|---|---|
| # | Coordinates of eccDNA by NGS | GC% | Genic sequence | Repeat region | Microhomology present within in 20 bp of flanking regions | |
| | | | | | 5' | 3' |
| 1 | Chr1:245530078-245530685 | 37.9 | KIF26B | Partial LINE | TACA | AGAAAAGG |
| 2 | Chr16:83620011-83620259 | 59.3 | CDH13 | None | TGG | GGA |
| 3 | Chr12:117100381-117100661 | 54.8 | TESC-AS1 | None | TT | TGC |
| 4 | Chr6:43488800-43490545 | 41.4 | TJAP1 | None | CA | TGG |
| 5 | Chr14: 69346689-69347123 | 57.8 | GALNT16 | None | ACA | TCT |
| 6 | Chr11: 100650414-100650783 | 48.9 | None | Partial SINE | TA | TG |
| 7 | Chr1:245530303-245530648 | 40.7 | KIF26B | None | AGC | AT |
| 8 | Chr19:49091220-49091256 | 56.1 | SNRNP70 | None | CC | CT |

B

| DT40 Cell Line EccDNA Candidates | | | | | | |
|---|---|---|---|---|---|---|
| # | Coordinates of eccDNA by NGS | GC% | Genic sequence | Repeat region | Microhomology present within in 20 bp of flanking regions | |
| | | | | | 5' | 3' |
| 1 | Chr11: 2300736 - 2300969 | 42.7 | CR406291 | None | GA | GAT |
| 2 | Chr4: 1460467 - 1462710 | 53.3 | DR416172 | None | GC | CT |
| 3 | Chr3: 25576742 - 25577047 | 33 | BX932473 | None | AG | TA |
| 4 | Chr4: 82442326 - 82443624 | 36.6 | None | None | ATT | TCA |
| 5 | Chr2: 69137047-69137904 | 35.4 | None | None | TAT | TC |
| 6 | Chr8: 13132654 - 13133032 | 44.6 | None | None | GGCAC | GA |
| 7 | Chr2: 20383321 - 20383744 | 43.9 | X58519 | None | ACC | AG |
| 8 | Chr1: 30701201 - 30701303 | 37.9 | None | None | CAG | TA |

**Supplemental Table 3: Inverse PCR primers to detect eccDNA formed from near a CRISPR-Cas9 induced double strand break on Chr22 and Chr12.** Sequences of outward facing primers, size of inverse PCR amplicon and sequence of the amplicon with the two parts that are joined to form the circle are given. The two parts are indicated by bold and regular font. When the primers are spaced apart, the entire eccDNA sequence is not amplified, and the PCR amplicon length is smaller than the entire length of the eccDNA molecule from which it was amplified.
We believe that the eccDNA does not contain the genomic region exactly up to the DSB because of (1) resection of the DNA after the DSB or (2) reconfiguration of the DNA during the repair.

| EccDNA Induced by DSB: OF Primers and Amplicon Sequences | | | | | |
|---|---|---|---|---|---|
| # | UF Primer sequence | DF Primer Sequence | Coordinates of eccDNA by NGS | Size of Amplicon (bp) | Sequence of OF primer PCR amplicon |
| 1 | CCCAGTGCACCCCAACAAAGTGTTGAT | AGGGGAGGTGAGGGGTGAGTGAG | Chr22: 18,623,231-18,624,741 | 1225 | **CTGGGCTGGGACCGTGGTCTGGATGCCAGGGAGTGGAAAGGAGCTGAAGGAAC**<br>**TCTCGCAGGCGGGCAGGCCTCCCAAGGGATTGGTTCAGCCAGAGAGGCAGGAC**<br>**ACGGACGGGCCAAGTCTCGGTGTGCATAGATTAGCGCTGGGGAAAGTCCCCGTG**<br>**GGGGCAGGCTGTGGGCAGGGAGGAGGGCTCACCAGCCAGCCAAGTGCCCGCAC**<br>**GAGGGAAGGAGTGGCCATCGTCCAGTGAGATGGTGAAGGGGACGTGGCCGCTC**<br>**TCATAGAGCAGGGGCGACACGAAGTGCACTTGCCCGGAGG**GTTTCACCACATTG<br>GCCAGGATGGTCTTGATCTCCTGACATCATCGTGATCCGCCCGCCTTGGCCTCCCA<br>AAGTGCTGGGATTACAGGCGTGAGCCACCGTAACCAGCCTGCCCTGCGTAATTTTT<br>GTTGTCGATGATCCAAAGTGCACCCACTCGCCATCTCTGGTAGCTGGGAGGAGCAG<br>GTTTTTGGGTGAAGGTGAAAGAGCCGGAGTTGGGGATGTGTGTGGCTGGGGGGTA<br>CAGGTGCGACCACTTTGCAGTCCACTCCTGTGAGTAGGGCATTCCTGAGGGAGAG<br>AAGGCCTGGTTAGCCCTGGGGCTCCTTCAAGGCCTGGCCCCAGGCCTTGATACCT<br>AGAGCAGCCTGACACATGTGTGGATGGACTGCCAGGCGGTCCCCTTAGGAGGGGC<br>TGTTCTAGCAGGGACCCTGGCTTGGGTGACCACCACCTCCTGCCACACTCCAGGA<br>TCCCTGTCCCTGCTGGAAGCCGGGCCCAGGGCCATGCCAGCCGTGAGGGGTTCC<br>TCATGAGCCTCCTCCCCGGGACCCAGGCCCCATGCAGCTGCCCCCTCCCACCTGC<br>CTGACCCCCCTGGCCTGCACTTCCCTCCATCCGAATCCTCAATCTCCGCCTGGAAT<br>TCCCTTCCCTCCTCAACCCCACCTGGAAACTCTCCTCCTCCTCCCACACAGCTGGG<br>ACTCCCTTTCTCCCACCTCTGGGGGCCCAAAGCTCTGGATGCCACCCTAGCCCTCA<br>GCTGGCCTCACCTGTCTCCTCATAGCCCCACAGCTTGATGGTGTCTGTGTGGGCAG<br>CGCCTCAACGTGCCAGGTCAGGCTGAGGTTGCCTGAGGTGTTGGCGGTGCCGTA<br>GTATTGCCAATGCATCTCGTTCATCAACTCACTCTTCTCCTTCATCAACACTTTGTTG<br>GGGTGCACTGGGCAGGG |
| 2 | CCAGTGCCTATCCACTGAGCTCCG | GTCCCAGCCCAGAGTGAGTGGG | Chr22: 18,624,003-18,624,948 | 680 | **TGCTCTCCTGTGGCATGCAAACCTGTGGGCCTGGTGGTGGTAGGAGGTGCTGGG**<br>**CCACTGCAGGCTGAGCATCTCGACAGTCTGGGCCCCCAACCTCCCTGGGGAACC**<br>**CATCCAACCAGCTGTCCTGTGCCCTCCTGCCTGTCCTGGGTCTGAGGGGCTCCAT**<br>**TCAGCCTGCCAGTGGGTCTGAGCCCTCAGCATGCACAGGGATGGGGGCGTGGGG**<br>**CCCCAGTGCCCGGCCTCCTGGGCCTCCCAACCTGCAGATCAC**CCCTCGGCCATG<br>TGGACTCCTCCGGGCAAGTGCACTTCGTGTCGCCCCTGCTCTATGAGAGCGGCCA<br>CGTCCCCTTCACCATCTCACTGGACGATGGCCACTCCTTCCCTCGTGCGGGCACTT<br>GGCTGGCTGGTGAGCCCTCCTCCCTGCCCACAGCCTGCCCCCACGGGGACTTTC<br>CCCAGCGCTAATCTATGCACACCGAGACTTGGCCCGTCCGTGTCCTGCCTCTCTGG<br>CTGAACCAATCCCTTGGGAGGCCTGCCCGCCTGCGAGAGTTCCTTCAGCTCCTTTC<br>CACTCCCTGGCATCCAGACCACGGTCCCAGCCCAGAGTGAGTGGGAGCTGCAGG<br>GGTCCCTAGAGAGGTGGGCCAGTGCCTATCCACTGAGCTCCGCCACACCAGGGCA<br>GGGGAGAAGCCAGGTGG |
| 3 | CCCTCGGCCATGTGGACTCCTC | GCTCTCCTGTGGCATGCAAACCTGT | Chr22: 18624345-18625024 | 946 | **GGGAGGTGAGGGGTGAGTGAGGGCTGGGCCCAGGGAAGCAGGGAGGAGGCCC**<br>**TGAGGCTTCAAGGGCTGTGGGAGCCTCCAGGGCCAATGCATGGCAGCAGCTGGG**<br>**TGATGGTCTCTCCGGCCCTCATGGCAGAAAAGGACGCTGAGGCCGGAGAGGCAG**<br>**GGGCCTGCCCTCCCTCCACTGCCCACGCCTCTAGCCTCCACCTGGCTTCTCCCCT**<br>**GCCCTGGTGTGGCGGAGCTCAGTGGATAGGCACTGGCCCACCTCTCTAGGGACC**<br>**CCTGCAGCTCCCACTCACTCTGGGCTGGGACCGTGGTCTGGATGCCAGGGAGTG**<br>**GAAAGGAGCTGAAGGAACTCTCGCAGGCGGGCAGGCCTCCCAAGGGATTGGTTC**<br>**AGCCAGAGAGGCAGGACACGGACGGGCCAAGTCTCGGTGTGCATAGATTAGCGC**<br>**TGGGGAAAGTCCCCGTGGGGGCAGGCTGTGGGCAGGGAGGAGGGCTCACCAGC**<br>**CAGCCAAGTGCCCGCACGAGGGAAGGAGTGGCCATCGTCCAGTGAGATGGTGAA**<br>**GGGGACGTGGCCGCTCTCATAGAGCAGGGGCGACACGAAGTGCACTTGCCCGG**<br>**AGGAGTCCACATGGCCGAGGGTCTGGATGCTCTCCTGTGGCATGCAAACCTGTG**<br>**GGCCTGGTGGTGGTAGGAGGTGCTGGGCCACTGCAGGCTGAGCATCTCGACAGT**<br>**CTGGGCCCCCAACCTCCCTGGGGAACCCATCCAACCAGCTGTCCTGTGCCCTCC**<br>**TGCCTGTCCTGGGTCTGAGGGGCTCCATTCAGCCTGCCAGTGGGTCTG**TGGTGTC<br>TGTGTGGGCAGCGCCTCAACGTGCCAGGTCAGGCTGAGGTTGCCTGAGGTGTTG<br>GCGGTGCCGTAGTATTGCCAATGCATCTCGTTCATCAACTCACTCTTCTCCTTCATC<br>AACACTTTGTTGGGGTGCACTGGGCAGGG |
| 4 | ATAGCGCCCACCAGCCTCTG | CCTGGAGAGGCACACAGCCA | Chr12:117100381-117100661 | 269 | **ATAGCGCCCACCAGCCTCTGTGGCCATGTGCTGTGTCCACGCCAGGCCCTCTGGT**<br>**GGACATCTCATTAACTTATCCTTATTTTAAGGAGGAGGAATCACATTACACAGGCAA**<br>**CAGCATCGGCCCCAGTCACCCCAACTGGATGGAGA**AGACTAACGTTCAGAACAGA<br>CCTGATACCTGACGGTCATTGATGGTGGTGTCACTAAGCACTGCAGAGCCTTCAGG<br>AGAGCCTTGACCTTGATGTTCTGCCCTGGCTGTGTGCCTCTCCAGG |

**Supplemental Table 4:**
**The comparison of abundance of endogenous eccDNA from hotspots (bottom 8 rows) with eccDNA stimulated by DSB induction (rows labeled Chr22 gRNA).** The level of eccDNA stimulated by the directly targeted DSB at Chr 22 was not as high as the most abundant endogenous eccDNA from the hotspots 1 and 2, but were comparable to the eccDNAs from the other hotspots.

| | EccDNA category | Average abundance of eccDNA relative to mtDNA | Standard error of eccDNA levels relative to mtDNA |
|---|---|---|---|
| Control gRNA | DSB proximal eccDNA 1 | 2.72E-05 | 8.19E-06 |
| | DSB proximal eccDNA 2 | 4.63E-08 | 2.01E-08 |
| | DSB proximal eccDNA 3 | 1.55E-08 | 4.84E-09 |
| Chr22 gRNA | DSB proximal eccDNA 1 | 4.78E-05 | 2.06E-05 |
| | DSB proximal eccDNA 2 | 2.00E-07 | 7.44E-08 |
| | DSB proximal eccDNA 3 | 5.97E-08 | 1.80E-08 |
| Endogeno us eccDNA | Candidate eccDNA 1 | 4.03E-02 | 1.18E-02 |
| | Candidate eccDNA 2 | 1.04E-02 | 3.50E-03 |
| | Candidate eccDNA 3 | 7.99E-07 | 3.86E-07 |
| | Candidate eccDNA 4 | 4.69E-07 | 3.23E-07 |
| | Candidate eccDNA 5 | 4.02E-07 | 1.69E-07 |
| | Candidate eccDNA 6 | 9.05E-07 | 3.91E-07 |
| | Candidate eccDNA 7 | 4.79E-07 | 2.70E-07 |
| | Candidate eccDNA 8 | 2.30E-06 | 8.33E-07 |

**Supplemental Table 5: Induced DSB experimental design**: The characteristics of the (A) genomic region where DSB was induced, (B) sgRNA target site and of the eccDNAs detected, (C) eccDNA amplified by outward facing primers, (D) DNaseI sensitivity.

A

| Coordinates of region probed by outward facing primers | GC% | Chromatin accessibility | Repeat regions |
|---|---|---|---|
| Chr22: 18,623,992-18,634,095 | 60 | DNaseI Sensitive | Partial SINEs LINEs |
| Chr12: 117100085-117100681 | 54 | DNaseI Sensitive | Partial SINE |

B

| Coordinates of gRNA homology | GC% | Chromatin accessibility | Repeat regions |
|---|---|---|---|
| Chr22: 18,623,992-18,623,102 | 55 | DNaseI Sensitive | Partial SINE |
| Chr12: 117100085-117100105 | 55 | DNaseI Sensitive | None |

C

| # | Coordinates of eccDNA | GC% | Microhomology | Repeat regions |
|---|---|---|---|---|
| 1 | Chr22: 18,623,231-18,624,741 | 63 | None | Partial SINE |
| 2 | Chr22: 18,624,003-18,624,948 | 66 | None | None |
| 3 | Chr22: 18624345-18625024 | 66 | None | None |
| 4 | Chr12:117100381-117100661 | 53 | TCT | None |

D

**DNaseI sensitivity of region probed by outward facing primers by ENCODE Project: ENCBS634AAA**

Chr22: 18621395-18626503 region

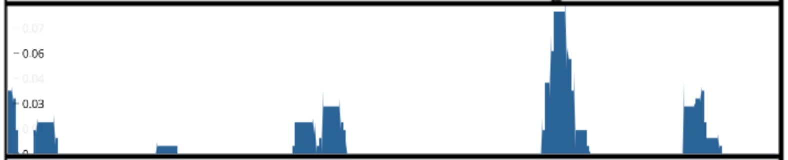


Chr12: 117069010-117162684 region

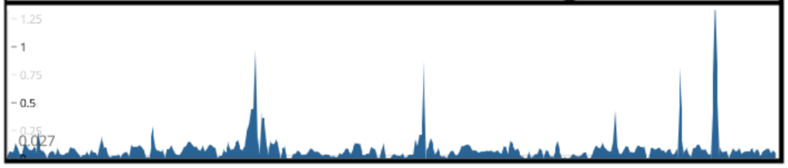
